# Host PGD_2_ acting on DP2 receptor attenuates *Schistosoma mansoni* infection-driven hepatic granulomatous fibrosis

**DOI:** 10.1101/2023.11.09.566508

**Authors:** Giovanna N. Pezzella-Ferreira, Camila R.R. Pão, Isaac Bellas, Tatiana Luna-Gomes, Valdirene S. Muniz, Ligia A. Paiva, Natalia R.T. Amorim, Claudio Canetti, Patricia T. Bozza, Bruno L. Diaz, Christianne Bandeira-Melo

## Abstract

Identifying new molecular therapies targeted at the severe hepatic fibrosis associated with the granulomatous immune response to *Schistosoma mansoni* infection is essential to reduce fibrosis-related morbidity/mortality in schistosomiasis. *In vitro* cell activation studies suggested the lipid molecule prostaglandin D_2_ (PGD_2_) as a potential pro-fibrotic candidate in schistosomal context, although corroboratory *in vivo* evidence is still lacking. Here, to investigate the role of PGD_2_ and its cognate receptor DP2 *in vivo*, impairment of PGD_2_ synthesis by HQL-79 (an inhibitor of the H-PGD synthase) or DP2 receptor inhibition by CAY10471 (a selective DP2 antagonist) were used against the fibrotic response of hepatic eosinophilic granulomas of *S. mansoni* infection in mice. Although studies have postulated PGD_2_ as a fibrogenic molecule, HQL-79 and CAY10471 amplified, rather than attenuated, the fibrotic response within schistosome hepatic granulomas. Both pharmacological strategies increased hepatic deposition of collagen fibers – an unexpected outcome accompanied by further elevation of hepatic levels of the pro-fibrotic cytokines TGF-β and IL-13 in infected animals. LTC_4_, one of the 5-lipoxygenase products described as potential anti-fibrotic mediators in the schistosomal liver, was reduced after HQL-79 and CAY10471 treatments. Moreover, LTC_4_ tissue levels were inversely correlated with collagen production in granulomatous livers. An ample body of data supports the role of *S. mansoni*-driven DP2-mediated activation of eosinophils as a key down-regulator of schistosomiasis-induced liver fibrosis, including: (i) HQL-79 and CAY10471 impaired systemic eosinophilia, drastically decreasing eosinophils within peritoneum and hepatic granulomas of infected animals; (ii) peritoneal eosinophils were identified as the only cells producing LTC_4_ *in* PGD_2_-mediated *S. mansoni*-induced infection; (iii) the magnitude of hepatic granulomatous eosinophilia positively correlates with *S. mansoni*-elicited hepatic content of cysteinyl leukotrienes, and (iv) isolated eosinophils from *S. mansoni*-induced hepatic granuloma synthesize LTC_4_ *in vitro* in a PGD_2_/DP2 dependent manner. So, our findings uncover that PGD_2_ by activating DP2 receptors stimulates cysLTs production by eosinophils, while endogenously down-regulates the hepatic fibrogenic process of *S. mansoni* granulomatous reaction – an *in vivo* protective function which demands caution in the future therapeutic attempts in targeting PGD_2_/DP2 in schistosomiasis.

**Author summary:** Accumulation of scar tissue (fibrosis)in the liver is the main cause of health problems associated with schistosomiasis even after the parasite is eliminated by treatment with anthelmintics. Previous experiments with isolated cells in culture have identified a potential role for a lipid mediator, PGD2, in promoting liver fibrosis leading to suggestions that PGD2 inhibition may be beneficial for the people infected with Schistosoma parasite. However, there was no direct evidence in an infection model to support these claims. Here, we described the effect of inhibiting the production or action of PGD2 in a mouse model of schistosomiasis. We identified the cell target and mechanism of action of PGD2’s participation in schistosomiasis. However, our data indicates that PGD2 protects the liver from fibrosis. Thus, inhibition of PGD2 action in patients infected with the Schistosoma parasite may aggravate the condition and promote faster liver failure and should not be pursued as a treatment option.

## Introduction

Often resulting in portal hypertension and liver failure, liver fibrosis is a collateral outcome of hepatic diseases of varying etiologies, such as schistosomiasis – a neglected tropical immunopathology, whose fibrosis-driven morbidity and mortality affects millions of people infected with *Schistosoma spp* parasites worldwide. Current anti-schistosomal chemotherapy effectively kills worms resolving the infection with only mild side effects. Yet, controversy persists about whether the present therapeutic approach successfully treats the schistosomiasis-induced liver fibrotic sequelae [1–3]

In *Schistosoma mansoni* (*S. mansoni*) infection-driven pathogenesis, hepatic fibrosis is a long-lasting feature of the excessive granulomatous inflammation built around liver-trapped eggs. The assembly of *S. mansoni*-induced hepatic fibrotic granulomas follows a switch from an initial type 1 to the pro-fibrotic type 2 immune response. And though protective at its onset, the large extensions occupied by the persistent fibrotic granulomatous tissue can culminate with disruption of the normal liver architecture and loss of its essential functions. Initiated by the egg deposition in the liver parenchyma and orchestrated by type 2 cytokines and intense eosinophilic inflammation, the hepatocellular response within schistosomal granulomatous space triggers complex cellular events – remarkably, the transdifferentiation of resident stellate cells into hepatic myofibroblasts [4]. Intra-granulomatous activation of stellate myofibroblast evokes prominent collagen synthesis and periovular deposition in the *S. mansoni*-infected liver parenchyma [5]. It is well-established that the granulomatous myofibroblast-ruled fibrogenic process is notably promoted by transforming growth factor-β1 (TGF-β1) and IL-13; indeed, the specific blockade of TGF-β1 and IL-13 reduces the schistosomal fibrosis [6,7]. Of note, besides their role in fibrosis as the pivotal collagen-synthesizing cells, *S. mansoni* infection-activated stellate myofibroblasts also appear to display additional immunomodulatory functions through the release of chemokines, cytokines, and bioactive lipid mediators, such as prostaglandin (PG)D_2_ [8]. PGD_2_ is a downstream metabolite of the arachidonic acid/cyclooxygenase (COX) enzymatic pathway, which is synthesized by the rate-limiting PGD synthase (PGDS) enzymes. In *S. mansoni* mammal hosts, in addition to the lipocalin-type PGDS highly expressed in the central nervous system, the hematopoietic PGD synthase (hPGDS) is the main terminal enzyme for PGD_2_ synthesis in peripheral tissues [9]. Alongside stellate myofibroblasts [8] various host cell types of the *S. mansoni*-driven inflammatory milieu express hPGDS and are recognized sources of PGD_2_ [9]. Moreover, eosinophils have been identified to generate PGD_2_ in an hPGDS-driven reaction within newly assembled cytoplasmic lipid bodies under inflammatory conditions. As a major cell component of schistosomiasis, eosinophils could also play a role in PGD_2_ generation within schistosomal granulomas [10,11].

Either autocrinally or paracrinally, PGD_2_ exerts most of its effects through the activation of two distinct 7-transmembrane G-protein coupled receptors, DP1 and DP2, co-expressed in cell surfaces. In contrast to the adenylate cyclase-driven inhibitory feature of DP1 activation, DP2 (formerly known as CRTH2) elicits the distinctive downstream signaling classically displayed by chemoattractant receptors. The resultant of concomitant activating both PGD_2_ receptors is a cell type-specific phenomenon [12,13]. In eosinophils, simultaneous activation of co-expressed DP1/DP2 classically triggers intracellular signaling pathways with opposing effects on chemotactic activity [14]. On the other hand, other eosinophil functions are a concerted activation of both receptors. For instance, for lipid body-compartmentalized leukotriene (LT)C_4_ synthesis, PGD_2_-induced concurrent DP1/DP2 activation converges toward triggering cooperative signaling and intracellular events [15,16,12]: DP1 brings about the assembly of new active lipid body compartments and DP2 activates LTC_4_ synthesizing machinery within newly formed lipid bodies [17,15].

The link between PGD_2_ and schistosomal immunopathology began 40 years ago with the observation that *S. mansoni* cercariae can generate PGD_2_ by themselves [18] and has advanced to unveil that (i) not only cercariae but also other *S. mansoni* life stages (remarkably eggs) produce PGD_2_ via the enzymatic activity of a schistosomal PGDS – the Sm28GST, a 28 kDa glutathione-S-transferase [19–21]; (ii) cercariae-derived PGD_2_ within infected skin elicits cutaneous immune evasion, decreasing the migratory capability of Langerhans cells towards draining lymph nodes [22]; and (iii) by employing a DP1 receptor-deficient mouse, activation of host DP1 receptor during *S. mansoni* infection was shown to be crucial for the establishment of early type 1 immune response (until 1 week) as well as late worm burden (7 weeks) in the liver [19]. Altogether, the data clearly show PGD2’s role in schistosomal immunopathogenesis and its potential as a novel adjuvant target for the current chemotherapy, focusing on quelling the deleterious impact of parasitic infection on the host.

Understanding the role of PGD_2_ after the host has been parasitized and before liver fibrosis has fully developed may generate medically relevant insight into disease management. Such studies are particularly relevant due to some divergent evidence for the role of PGD_2_ in fibrotic processes that affect other organs’ fibrogenesis. In renal settings, for instance, PGD_2_ arises as an anti-fibrogenic agent by suppressing the induction of fibrotic phenotype in cultured kidney cells [23,24]; in contrast, it also emerged as an interesting target for anti-fibrotic therapies of renal diseases since DP2 inhibition *in vivo* downregulates renal fibrosis in a chronic model of kidney inflammation [25]. In the lung, studies on non-asthmatic pulmonary fibrosis acknowledge PGD_2_ itself as a beneficial tool against fibrotic development [26–29], however, PGD_2_ displays clear pro-fibrogenic impact in the asthmatic lung [30,31]. Although simple extrapolation of PGD2’s role in liver fibrosis based on evidence gathered on different tissues may not be straightforward, it is reasonable to speculate that schistosomiasis-driven fibrosis may be promoted by PGD_2_, in a similar pro-fibrotic fashion as it is in asthma due to the shared type 2-governed eosinophilic environment.

Since PGD_2_ elusive function on fibrogenesis has not been properly validated using *in vivo* models of hepatic inflammation, the potential link between PGD_2_ and schistosomal hepatic fibrosis is mostly drawn from *in vitro* studies. While by employing DP1 deficient mice, no impact was observed in the fibrotic feature of hepatic granulomatous reactions around *S. mansoni* eggs [19], it has been observed that (i) host PGD_2_ synthesized by *S. mansoni*-driven stellate myofibroblasts in response to *in vitro* TGF-β autocrinally mediates myofibroblastic secretory activity [32]; (ii) *S. mansoni*-derived PGD_2_ acting via DP1 triggers *in vitro* eosinophil activation (lipid body biogenesis) [21] and (iii) induces 15-lypoxygenase-driven synthesis of eoxin C_4_, which in turn promotes TGF-β secretion [21]. Taken together with the pro-fibrotic role in type 2-biased asthma, these pieces of evidence build a case for a PGD_2_ role in promoting *S. mansoni*-driven fibrogenesis in hepatic granulomas. Moreover, such findings support recommendations of PGD_2_ as a possible target for anti-fibrotic therapies in schistosomiasis [33–35].

Our investigation of the role of PGD_2_ in schistosomal granulomas led to the discovery of endogenous counter-regulatory mechanisms required to limit hepatic fibrosis beyond those associated with DP1 receptor activation [19,36,37]. We have unveiled an endogenous PGD_2_/DP2 modulatory mechanism that reduces the granuloma-associated fibrotic reaction elicited by *S. mansoni* infection. Although missing pieces of information do not allow the drawing of a complete picture of the events responsible for PGD_2_-mediated inhibition of *S. mansoni* infection-elicited hepatic granulomatous fibrosis, our findings show that once oviposition starts, PGD_2_ (i) is produced within hepatic granuloma, (ii) activates DP2 receptors to mobilize blood eosinophils to hepatic granulomas, (iii) triggers LTC_4_ synthesis by granuloma-infiltrating eosinophils; while (iv) reduces hepatic production of fibrogenic TGF-β and IL-13; (iv) decreasing synthesis and deposition of collagen around granuloma-trapped eggs.

## Results and Discussion

### *S. mansoni* infection-induced hepatic granuloma formation triggers hPGDS-driven PGD_2_ synthesis

*In vitro* studies have suggested PGD_2_ as a potential molecular target for novel anti-fibrotic therapies in the hepatic granulomatous pathology of *S. mansoni* infection [8] [33,35]. To address such inference from *in vitro* data, here we first evaluate whether enhanced PGD_2_ production is elicited in an *in vivo* experimental model of schistosomiasis – C57BL/6 mice infected percutaneously with 60 cercariae of *S. mansoni* (**Fig 1A**). As shown in **Fig 1B**, the infection protocol triggers productive oviposition first detected in feces within 6 weeks, which peaks within 8 weeks post-infection (wpi), while no *S. mansoni* egg was found in feces as early as 3 weeks. Before oviposition (3 wpi), no increase in PGD_2_ synthesis was detected within peritoneal or hepatic compartments of *S. mansoni-*infected mice (**Fig 1C and D**). However, later at 8 wpi, PGD_2_ levels were elevated in both egg-containing granulomatous liver tissue, as well as, in peritoneal cavities of *S. mansoni*-infected mice (**Fig 1C and D**). Of note, we have previously demonstrated that parallel to the 8 weeks-related intense egg release in feces (**Fig 1B**) uncovered here, a type 2 immune response characterized by systemic eosinophilia is associated with the formation of egg-encasing hepatic eosinophilic granulomas with robust fibrosis around *S. mansoni* eggs [38,39]. Although regulatory roles for PGD_2_ in *S. mansoni*-induced type 2 immune response-biased pathology had been previously observed in functional studies employing the DP1 receptor-deficient mice [19], enhanced PGD_2_ production in the liver during schistosomiasis had not been demonstrated before.

**Fig 1.**
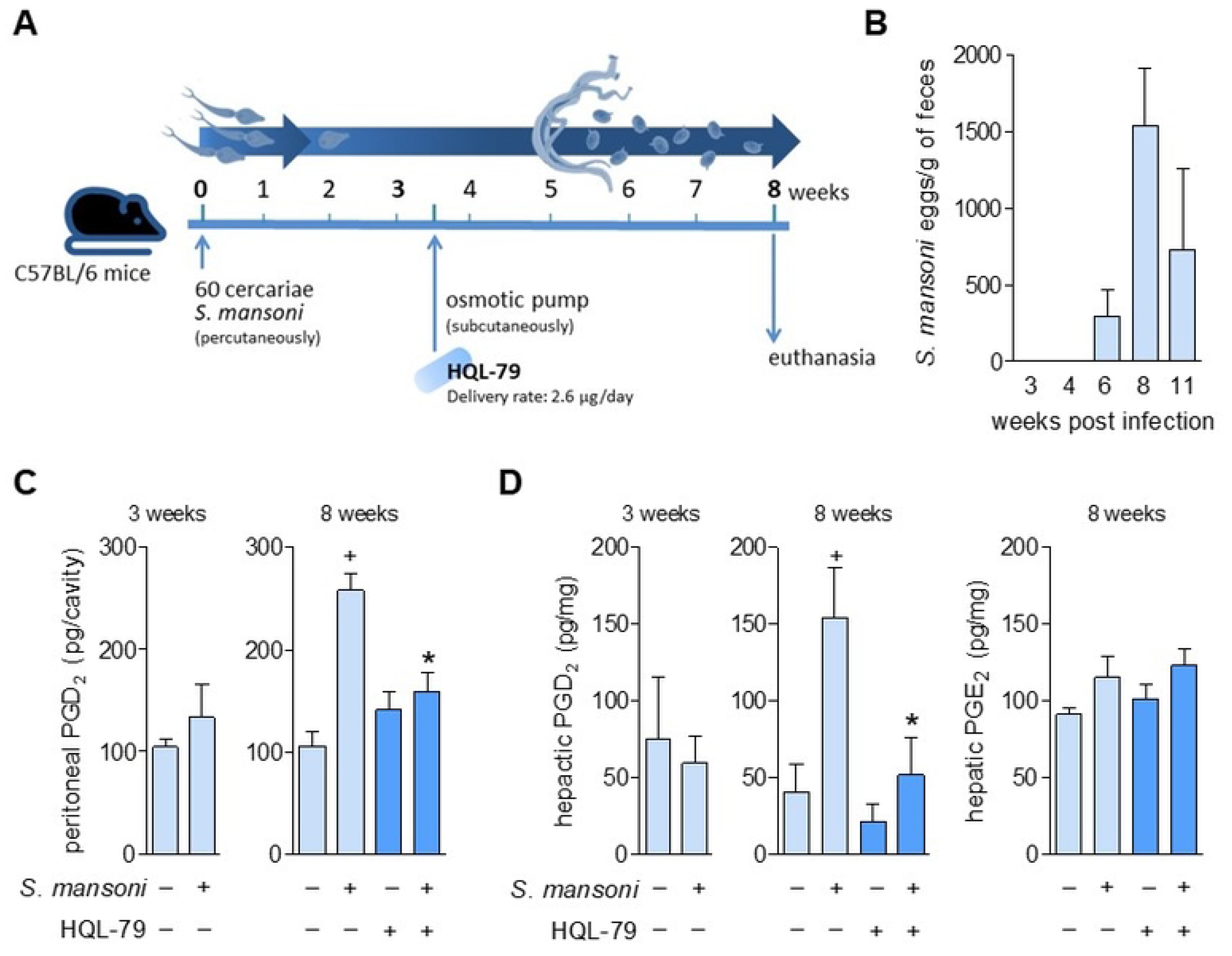
Characterization of a post-oviposition and host PGD synthase-mediated PGD_2_ synthesis during *S. mansoni* infection in mice. Scheme in **A** outlines the protocol of *S. mansoni* infection in mice achieved by active percutaneous penetration by 60 cercariae and the continuous treatment with HQL-79 (2.6 µg/day), which was delivered by subcutaneously implanted osmotic pumps at 3.5 weeks post infection (wpi) with *S. mansoni*. Peritoneal lavages and livers were collected 3 or 8 wpi. **B** shows a temporal kinetics of *S. mansoni* egg detection in feces. In **C**, PGD_2_ amounts detected by specific EIA kit in cell-free peritoneal fluids. In **D,** PGD_2_ or PGE_2_ levels in liver homogenates detected by specific EIA kits for each prostanoid. Values are expressed as the mean ± SEM from at least 5 animals *per* group (experiments were repeated at least once). ^+^*p* < 0.05 compared to non-infected control group. \**p* < 0.05 compared to infected non-treated group.

Administration of HQL-79 – a specific inhibitor of the hPGDS – effectively inhibited the host PGD_2_ synthesis induced by *in vivo S. mansoni* infection, thus confirming its adequacy as a pharmacological tool for our aims. As schematized in **Fig 1A**, to achieve systemic and persistent delivery of HQL-79 treatment during long-lasting *S. mansoni* infection, osmotic pumps containing HQL-79 (flow delivery rate of 2.65 µg/day; for about 30 days) were subcutaneously implanted at 3.5 wpi – therefore, before either oviposition (**Fig 1B**) or the onset of *S. mansoni* infection-induced increase of systemic PGD_2_ levels (**Fig 1C and D**). Validating the systemic reach of this administration strategy, as well the HQL-79 inhibitory specificity towards PGD_2_ synthesis, HQL-79 treatment did impair *S. mansoni* infection-induced PGD_2_ synthesis detected in both peritoneal wash and liver (**Fig 1C and D**), while did not alter hepatic tissue PGE_2_ levels (**Fig 1D**).

### Inhibition of PGD_2_ synthesis or DP2 antagonism aggravates fibrosis of *S. mansoni*-induced hepatic granulomas

The proposed pro-fibrogenic function of endogenous PGD_2_ in hepatic granulomatous response to *S. mansoni* infection [33,35] was investigated by employing two complementary pharmacological strategies, including either (i) the proven *in vivo* PGD_2_ synthesis inhibition by HQL-79 treatment **(Fig 1C** and **1D**); or (ii) the blockade of PGD_2_ receptor DP2 by a selective antagonist, CAY10471, also delivered by osmotic pump system (flow rate of 1.75 µg/day; implanted at 3.5 wpi). As shown in **Fig 2**, *S. mansoni* infection triggered an intense periovular granulomatous response in livers, presenting enhanced levels of TGF-β and IL-13, as well as locally deposited collagen around *S. mansoni* eggs entrapped in hepatic granulomas. In contrast to the hypothesis of a pro-fibrogenic role of PGD_2_ in schistosomiasis derived from *in vitro* studies, in our experiments, both PGD_2_-targeting treatments aggravated, rather than attenuated, the *S. mansoni*-induced fibrotic reaction in the granulomatous liver. Such unexpected outcome was evidenced by a variety of fibrosis-related hepatic parameters, which were further enhanced by HQL-79 and CAY10471 treatments (**Fig 2**), including *S. mansoni* infection-induced: (i) increased hepatic collagen synthesis (as assessed by total hydroxyproline liver content) (**Figs 2A and 2B**); (ii) deposition of collagen fibers around *S. mansoni* eggs in hepatic granulomas (**Fig 2A and 2B**); (iii) hepatic production of the key pro-fibrogenic cytokine TGF-β (**Figs 2C and 2D**); and (iv) hepatic levels of the prototypical type 2 immune response cytokine IL-13 which is known to display potent fibrogenic activity in liver as well (**Fig 2C and 2D**). Considering mechanistic aspects, either HQL-79 (**Fig 2A**) or CAY10471 (**Fig 2B**) seemed to worsen overall hepatic *S. mansoni* fibrosis by specifically increasing localized collagen synthesis within granulomas which were already in development; since both treatments: (i) failed to modify the overall number of egg-encasing granulomas found 8 wpi within *S. mansoni*-infected livers, while (ii) promoted evident expansion of fibrotic area of individual *S. mansoni*-driven granulomas (**Figs 2A and 2B**). Notably, these findings also indicate that DP2 receptors activation by endogenous PGD_2_ – that occurs between 3.5 and 8 wpi – does not play key roles in controlling total egg production or the hepatic burden of parasites. It is important to highlight that our experimental design does not allow general assumptions on whether DP2 activation has endogenous roles in the initial cutaneous phase, oviposition-onset, or granuloma assembly. It is noteworthy that, inversely, DP1 activation by PGD_2_ during *S. mansoni* infection appears to negatively impact early parasite survival and the successful establishment of infection [19]; a caveat being that DP1 role on *S. mansoni* parasitism is derived from studies with genetically modified mice [19]. Therefore, DP1 function in an ongoing infection (*i.e.*, after the conclusion of the cutaneous phase) still needs to be pharmacologically addressed in schistosomiasis. Furthermore, based on classical opposing effects typically displayed by DP1 *versus* DP2 activation [40,41], selective antagonism of DP1 receptor could even trigger beneficial anti-fibrogenic outcomes.

**Fig 2.**
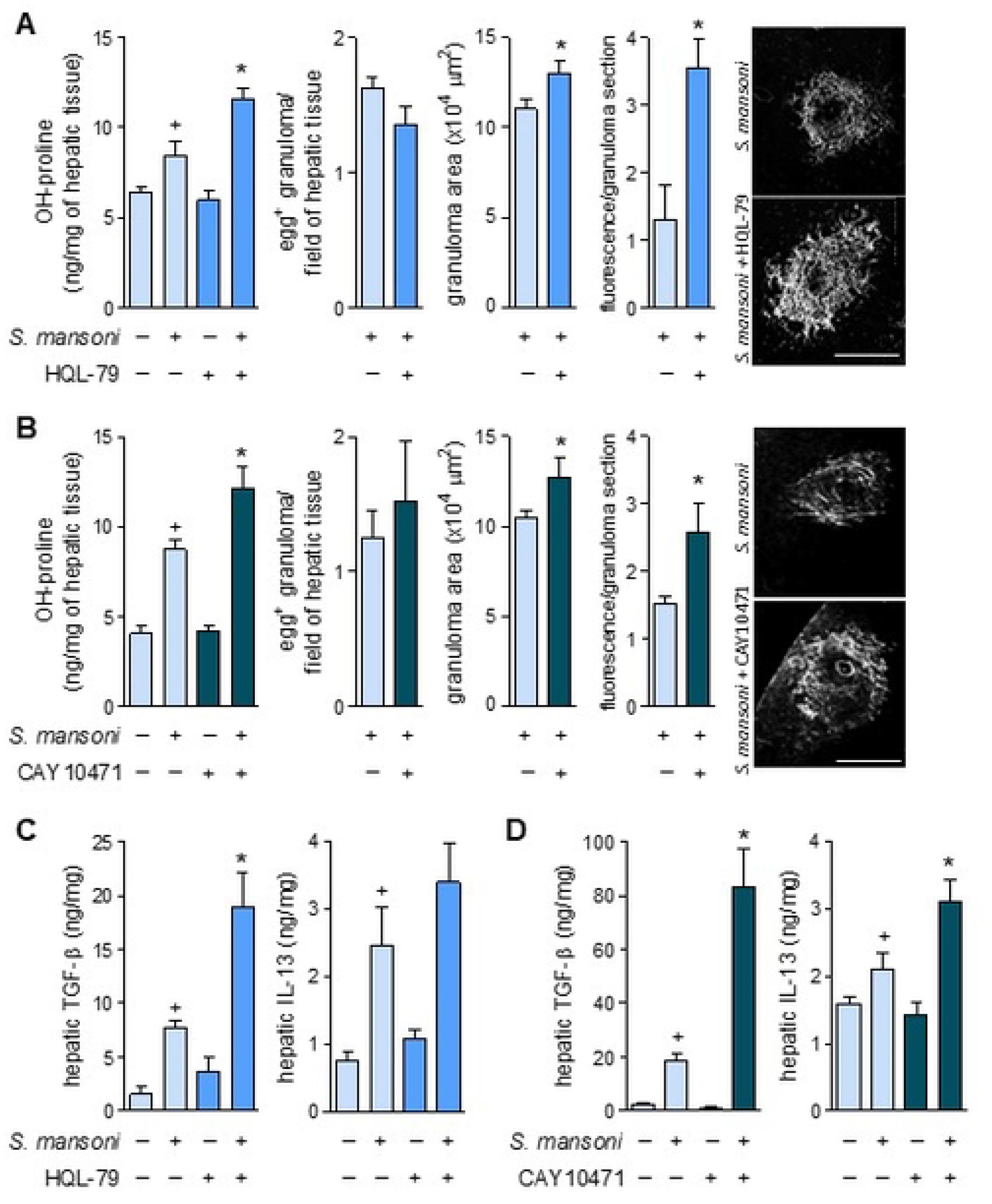
Inhibition of PGD_2_ synthesis by HQL-79 or selective antagonism of DP2 receptor by CAY10471 promote amplification of overall fibrotic reaction within *S. mansoni*-infected hepatic tissue. Continuous treatments (delivered by implanted osmotic pumps) with either HQL-79 (2.6 µg/day) or CAY10471 (1.7 µg/day) were initiated at 3.5 wpi with *S. mansoni*. Livers were collected 8 wpi. **A** and **B** show total amounts of hepatic hydroxyproline, total numbers of egg-encasing hepatic granulomas, and area measurements of individual hepatic granulomas. Images are representative collagen-related fluorescence in hepatic granuloma sections (confocal microscopy; PicroSirius red modified staining). **C** and **D** show levels of TGF-β and IL-13 detected by specific ELISA kits in liver homogenates of *S. mansoni*-infected mice. Values are expressed as the mean ± SEM from at least 5 animals *per* group from representative experiments for each drug studied. ^+^*p* < 0.05 compared to non-infected control group. \**p* < 0.05 compared to infected non-treated group.

The unexpected pro-fibrotic effect of both HQL-79 and CAY10471 treatments was accompanied by a further increase of the infection-elicited hepatic levels of the pro-fibrogenic factors, TGF-β and IL-13 (**Fig 2C** and **2D**). So, our findings also indicate that the mechanisms involved in the PGD_2_/DP2-driven down-regulatory function may depend on counterbalancing *S. mansoni*-induced hepatic production of TGF-β and IL-13 *in vivo*. Strikingly, as already mentioned previously, *in vitro* studies showed a diametrically opposite pattern for PGD_2_ regulation of TGF-β or IL-13 secretion. For instance, PGD_2_ via DP2 receptor activation is known to trigger, rather than inhibit, IL-13 production from isolated ILC2 cells [42]. Regarding *S. mansoni* infection, we have demonstrated *in vitro* that (i) PGD_2_ participates in *S. mansoni*-induced eoxin C_4_-driven TGF-β release from eosinophils [21] and (ii) PGD_2_ released by hepatic stellate cells isolated from *S. mansoni* granulomas autocrinally contributes to TGF-β-induced activation of these collagen secretory cells [32]. The *in vitro versus in vivo* discrepancies unveiled here for the role of PGD_2_ in *S. mansoni*-driven fibrotic process are likely due to the single cell type feature of *in vitro* assays, which lacks the sequential cellular activities performed by multiple cell types (*vide infra*) working in the more complex *in vivo* setting of hepatic granulomatous reaction. Of note, besides their significant role in the establishment of fibrosis [43,44], these *S. mansoni*-driven hepatic myofibroblasts are also recognized to contribute to local eosinophilia of hepatic periovular granuloma [8].

### Endogenous PGD_2_ mediates *S. mansoni* infection-elicited eosinophilic inflammation

Eosinophils have recently emerged as protective agents in hepatic tissue, acting towards reinstating liver homeostasis after diverse types of injuries [45,46]. Hepatic granulomatous eosinophilia is an undisputable hallmark of *S. mansoni* infection, yet these cells are considered at most minor immunomodulators of the protective type 2 immune response or even simple bystanders of the pathogenesis [47–49]. On the other hand, it is well-established that eosinophils represent key players in PGD_2_-regulated inflammatory responses since they function as both cell sources and targets of PGD_2_ [15] [40,11]. With an experimental design not aiming to re-examine eosinophil role in *S. mansoni*-driven immunopathology, here HQL-79 effect on *S. mansoni* infection-elicited hepatic eosinophilic reaction was examined seeking novel insights on the molecular/cellular mechanisms involved in the unexpected anti-fibrogenic effect of PGD_2_.

Reproducing clinical schistosomiasis, the mouse experimental model of *S. mansoni* infection used here exhibits, within 8 wpi, a marked systemic eosinophilia characterized by elevated numbers of circulating eosinophils, as well as infiltrating eosinophils found in both peritoneal cavities and in egg-encasing hepatic granulomas (**Fig 3**). Eosinophil recruitment from the circulation into *S. mansoni* infection-elicited inflammatory sites is known to be mediated by a variety of parasite- and host-derived chemoattractant molecules, especially host CCL11 [50]. Based on the potent eosinophilotactic activity of PGD_2_ [14,51], we analyzed whether endogenous PGD_2_ would also contribute to *S. mansoni* infection-induced eosinophilic reaction. As shown in **Fig 3**, by systemically inhibiting host PGD_2_ synthesis, HQL-79 was capable to impair the establishment of *S. mansoni*-driven eosinophilia, decreasing blood eosinophil availability (**Fig 3A, left panel**), eosinophil migration to peritoneal cavities (**Fig 3A, right panel**) and eosinophil influx into *S. mansoni*-induced hepatic granulomas (**Fig 3B, left and middle panels**). Hence, our findings boost the status of hPGDS-derived PGD_2_ to an important host-derived mediator of *S. mansoni* infection-elicited systemic eosinophilia. The clear inhibitory effect of HQL-79 towards the eosinophilic feature of *S. mansoni*-driven inflammatory response indirectly uncovers at least three mechanistic aspects of PGD_2_-mediated regulation of the fibrogenic process of schistosomal hepatic granulomas.

**Fig 3.**
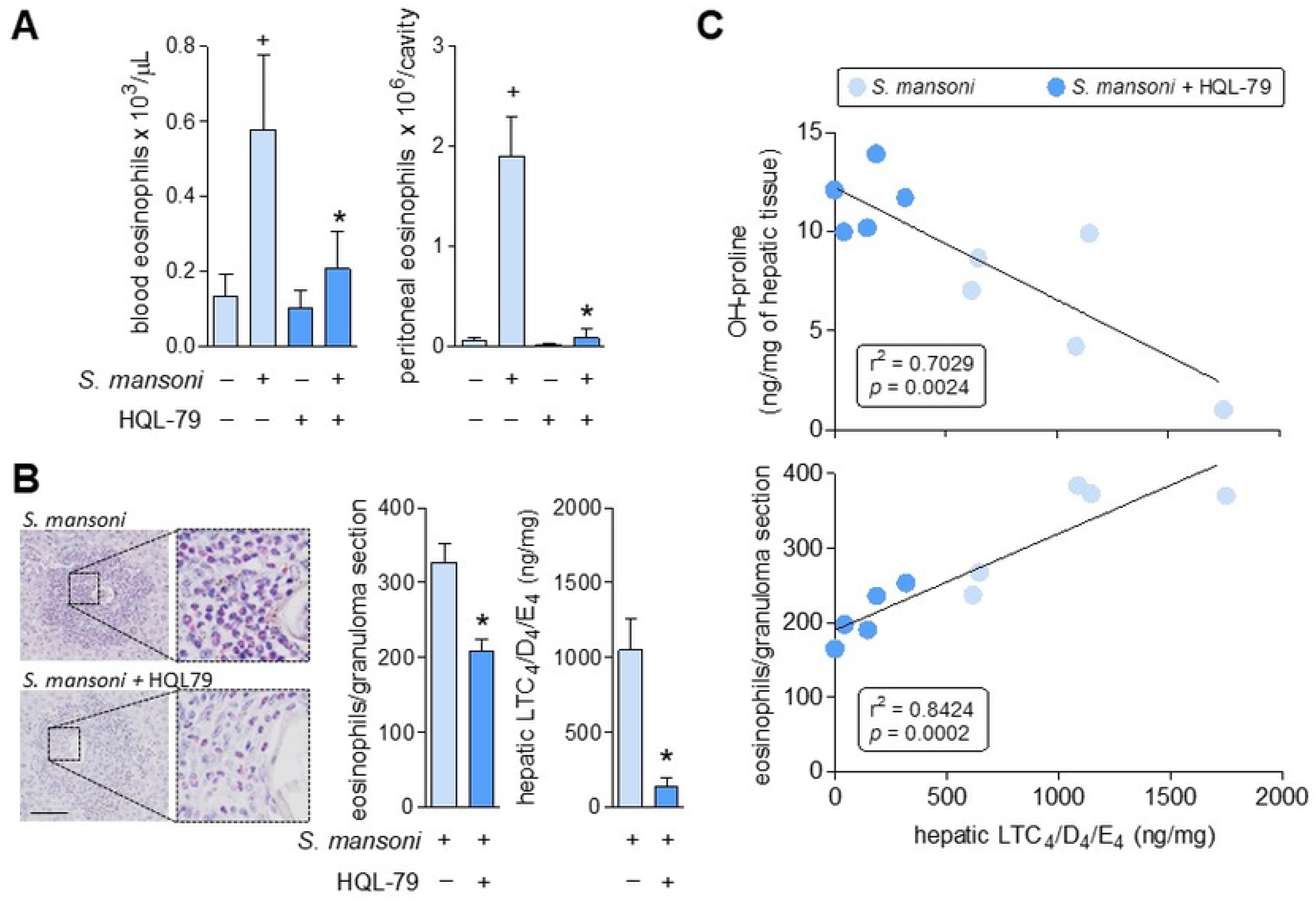
HQL-79 inhibits eosinophilic reaction and cysLTs production induced by *S. mansoni* infection. **A** shows eosinophil numbers found in either peripheral blood or peritoneal compartments 8 wpi with *S. mansoni*. **B** highlights the eosinophilic feature of hepatic granulomas (stained by Sirius red), showing representative images and enumeration of granuloma-infiltrating eosinophils as well as the amounts of cysLTs (LTC_4_/D_4_/E_4_) detected by specific ELISA kits in liver homogenates. **C** displays linear regression curves illustrating the relationships between cysLTS levels and either collagen synthesis (top panel) or granulomatous eosinophilic reaction (bottom panel) found in hepatic tissues. Values are expressed as the mean ± SEM from at least 5 animals *per* group (experiment was repeated at least once). ^+^*p* < 0.05 compared to non-infected control group. \**p* < 0.05 compared to infected non-treated group. R-squared (r^2^) and *p* values for each linear regression are shown in each panel.

First, distinct from the eosinophilic reaction impairment observed in DP1 receptor-deficient model [19], HQL-79-unveiled PGD_2_ role in *S. mansoni* infection-driven eosinophilia does not correspond to an inevitable consequence of a large inhibition of *S. mansoni* parasitism or overall lack of subsequent type 2 immune response. The inhibition of PGD_2_ synthesis by HQL-79 diminished *S. mansoni* infection-triggered eosinophilic reaction, although the magnitude of parasitic burden (number of eggs trapped in hepatic tissue) was unaffected (**Fig 2A**) and associated with a clear type 2 immune environment. Indeed, besides the slightly elevated (although not statistically significant) hepatic IL-13 levels found under HQL-79 treatment (**Fig 2C**), the PGD_2_ synthesis inhibitor failed to change the increased serum levels of type 2 cytokines IL-5 and IL-13 (**S1A Fig**) as well *S. mansoni* infection-triggered eosinophil production at mouse bone marrows (**S1B Fig**), a known IL-5-driven phenomenon.

Second, although known PGD_2_ producers [10], at first glance granulomatous eosinophils did not seem to be the main host cell producers of PGD_2_ at hepatic inflammatory site during *S. mansoni* infection, since: (i) HQL-79-promoted reduction of PGD_2_ derails the establishment of hepatic granulomatous eosinophilia (**Fig 3B**), and (ii) *S. mansoni* infection-induced PGD_2_ synthesis (as analyzed at 6 wpi) precedes eosinophil appearance (not shown). Other hPGDS-expressing cell types which are structurally present at schistosomal granuloma before PGD_2_-mediated eosinophil arrival at hepatic granulomatous tissue may be the main responsible for PGD_2_ synthesis. We have previously shown that the *S. mansoni*-driven hepatic stellate cells (granulomatous myofibroblasts) produce PGD_2_ in the schistosomiasis context in an HQL-79-sensitive manner [8]. Inasmuch as *S. mansoni*-driven myofibroblasts are also known to regulate eosinophilic reaction [32], we can hypothesize that hPGDS expressed by stellate cells are the target of HQL-79, which mediates the observed effects, including inhibition of hepatic eosinophilia and augmentation of fibrosis-related parameters. Nevertheless, the possibility that eosinophils within hepatic granulomas may contribute to the bulk of PGD_2_ detected by 8 wpi cannot be dismissed – further experiments are needed in order to decipher the role of stellate cells and eosinophils as possible sources of PGD_2_.

In this regard and thirdly, eosinophil does not seem to be the cell source of TGF-β or IL-13 in *S. mansoni*-infected livers. Even though eosinophils are known cell reservoirs of preformed TGF-β and IL-13 [47,52], HQL-79 decreased eosinophil presence (**Fig 3B**), whilst increasing the fibrogenic cytokines (**Fig 2C**) in *S. mansoni*-driven hepatic granulomas. Such inversed relationship indicates that, once in place within hepatic granulomas, eosinophils may suppress, rather than secrete TGF-β and IL-13 themselves or further their release by other granuloma cell types. At the granulomatous microenvironment, a paracrine downregulatory impact, likely orchestrated by secretory activities of PGD_2_-stimulated eosinophils, may indeed ensure the lower release of TGF-β and IL-13 by other cell sources.

### PGD_2_ elicits LTC_4_ production during *S. mansoni* infection: role of eosinophils as LTC_4_ synthesizing cells

The lipid molecules LTC_4_, LTD_4,_ and LTE_4_, collectively known as cysteinyl leukotrienes (cysLTs) are notoriously pro-fibrogenic mediators in a variety of lung conditions [53–55]. Therefore, based on pulmonary studies, it is anticipated pro-fibrogenic activities for cysLTs in other organs. However, for hepatic fibrosis of any etiology, the role of these molecules is yet uncharacterized; a notable exception is a study performed by Toffoli and coworkers (2006) showing that 5-LO-derived metabolites seem to act as suppressing signals of hepatic granulomatous fibrosis in *S. mansoni* infection [56,57]. Here, HQL-79 treatment decreased total amounts of cysLTs found in *S. mansoni-*infected hepatic tissue (**Fig 3B**). Concurring with a cause/effect relationship, and therefore a likely anti-fibrogenic function for these lipid molecules in schistosomal granuloma of infected livers, **Fig 3C** (top panel) shows a significative inverse correlation (r^2^ = 0.7029; *p* ≤ 0.05) between hepatic levels of cysLTs and the fibrosis marker hydroxyproline. However, direct and systematic verification of whether hepatic cysLTs down-modulate *S. mansoni*-driven granulomatous fibrosis is still pending.

*S*. *mansoni* infection-induced egg-entrapping hepatic granulomas are comprised of various functionally active cells, including hepatic stellate myofibroblasts, mast cells, macrophages, and eosinophils [50,39,58]. In addition to *S. mansoni* eggs themselves, all these granuloma-associated cells present the ability to synthesize LTC_4_ [59]. However, cysLTs production is not a ubiquitous cellular activity, it is rather a highly regulated, stimulus-specific, and cell-restricted phenomenon dependent on, for instance, cellular expression and proper intracellular localization of the limiting LTC_4_ synthase enzyme. Eosinophils express the entire LTC_4_ synthesizing enzymatic machinery [60,61], which is promptly activated and compartmentalized within cytoplasmic lipid bodies under PGD_2_ stimulation [15]. Therefore, at this point, we investigated that (i) *S. mansoni* infection-induced hepatic LTC_4_ synthesis may take place within PGD_2_-stimulated eosinophils infiltrating hepatic granuloma (*vide infra*); and (ii) the reduction of hepatic cysLTs observed under HQL-79 may be in part due to the decreased numbers of LTC_4_-synthesizing eosinophils within hepatic granulomas. Both assumptions are strongly supported by the significative positive correlation (r^2^ = 0.8424; *p* ≤ 0.05) between the magnitude of eosinophil presence and the levels of LTC_4_ found in each granuloma-enriched hepatic tissue of *S. mansoni* infected mice, treated or not with HQL-79 (**Fig 3C**; bottom panel). Altogether, eosinophils may not represent the main source of PGD_2_ or secrete TGF-β and IL-13 within schistosomal hepatic granulomas, but they may synthesize/release LTC_4_ under *in situ* stimulation by granulomatous PGD_2_ and be responsible for the increased hepatic levels of cysLTs.

### Production of LTC_4_ by *S. mansoni* infection-driven eosinophils upon PGD_2_ activation of DP2 receptors

Besides its eosinophilotactic activity, DP2 receptor is a required element for the successful induction of LTC_4_ synthesis by PGD_2_ in eosinophils – a cooperative intracellular process that (i) demands simultaneous activation of both DP1 and DP2 receptors, (ii) takes place within the discrete cytoplasmic lipid body organelles, and (iii) is triggered in both *in vitro*, as well as *in vivo* eosinophilic settings [11,12,62,63]. As shown in **Fig 4** and exactly like the inhibition of PGD_2_ synthesis by HQL-79, the selective antagonism of DP2 receptor by CAY10471 did decrease the number of infiltrating eosinophils found 8 wpi within the peritoneal space of *S. mansoni*-infected mice (**Fig 4A**). Infiltrating peritoneal eosinophils of non-treated infected mice displayed high cytoplasmic numbers of cytoplasmic lipid bodies (**Fig 4B**) as well as showed intracellular immunolabelling for newly synthesized LTC_4_ (**Fig 4C and D**). Antagonism of DP2 by CAY10471 (i) diminished the cytoplasmic content of lipid bodies within infiltrating peritoneal eosinophils (**Fig 4B**), (ii) shut down the ability of some peritoneal infiltrating eosinophils to synthesize LTC_4_ (**Fig 4C and D**), and therefore (iii) partially impaired the overall peritoneal production of cysLTs (**Fig 4E**) found 8 wpi with *S. mansoni*. In CAY10471-treated mice, residual peritoneal eosinophilia is still detected (**Fig 4A**) exhibiting a reduced (38.6 ± 6 % of inhibition; n = 5, *p* ≤ 0.05) LTC_4_ synthesizing ability (**Fig 4D and D**). Together these findings identify eosinophils as, at least, one of the cell types actively participating in *in vivo* PGD_2_/DP2 receptor-driven stimulation of LTC_4_ synthesis during *S. mansoni* infection. Moreover, the data indicate that the CAY10471-evoked whole decrease of peritoneal cysLTs levels (**Fig 4E**) may depend on an eosinophil-related three-component mechanism: (i) *in situ* partial loss of a LTC_4_ synthesizing cell population (the peritoneal eosinophils); (ii) decreased availability of intracellular LTC_4_ synthesizing compartments within DP2 activated eosinophils (lipid bodies); resulting in (iii) an impaired LTC_4_ synthesizing capability among remaining infiltrating eosinophils found in the peritoneal cavity.

**Fig 4.**
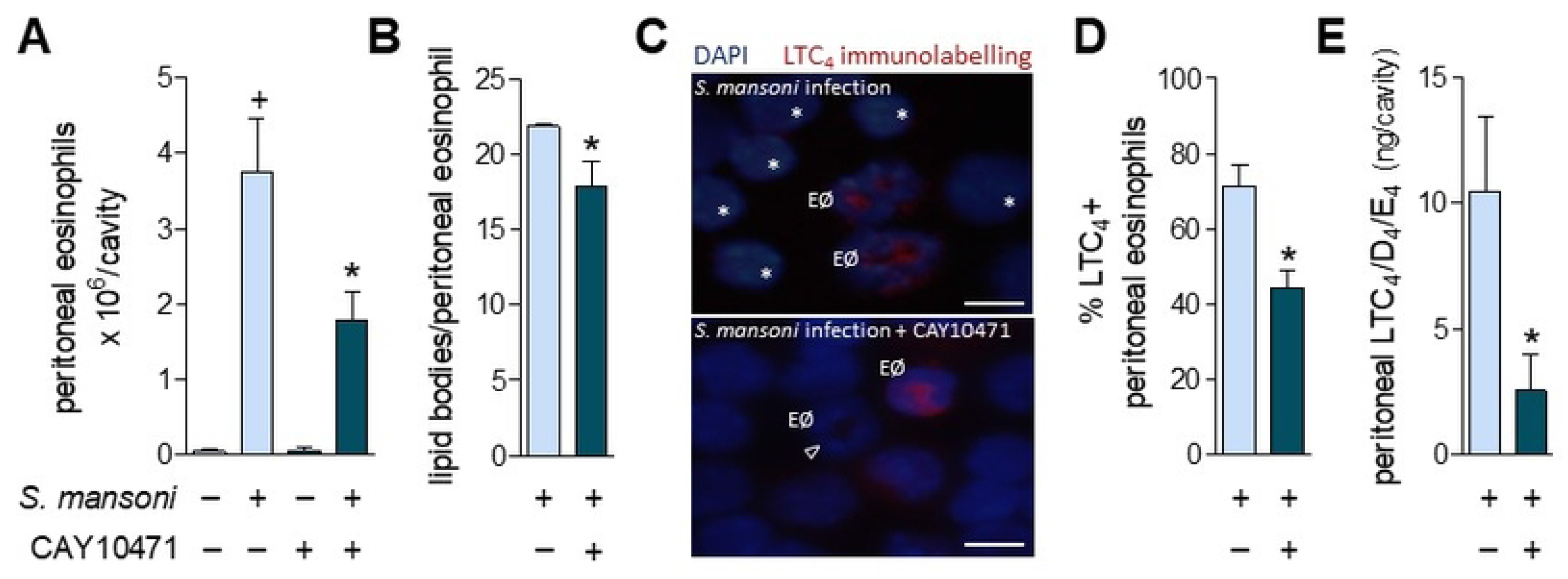
*S. mansoni* infection induces a PGD_2_/DP2-mediated cysLTs production by infiltrating peritoneal eosinophils. **A** shows peritoneal eosinophil counts while **B** displays the numbers of cytoplasmic lipid body organelles found into peritoneal eosinophils 8 wpi with *S. mansoni*. **C** displays representative images of EicosaCell preparations showing intracellular immunolabelled LTC_4_ (in red; LTC4^+^ eosinophils). Cellular nuclei are labelled with DAPI (in blue); bar = 10 µm; “EØ” identifies eosinophils among peritoneal cellular population; arrowhead indicates a LTC_4_^-^ eosinophil (remaining LTC_4_^-^ cells are mononuclear cells indicated by asterisks). In **D**, the percentage of peritoneal eosinophil population exhibiting cytoplasmic red immunostaining for LTC_4_ (LTC_4_^+^ eosinophils) is shown. **E** shows the total amounts of cysLTs (LTC_4_/D_4_/E_4_) detected by specific ELISA kit in peritoneal fluid supernatants. Values are expressed as the mean ± SEM from at least 5 animals *per* group (each experiment was repeated at least once). ^+^*p* < 0.05 compared to non-infected control group. \**p* < 0.05 compared to infected non-treated group.

As depicted in **Fig 4C** and **S2 Fig**, the peritoneal cell population found 8 wpi is not formed only by eosinophils, but by a mixed cell population of eosinophils and mononuclear cells. However, distinct from infiltrating eosinophils, the *S. mansoni* infection-elicited peritoneal mononuclear cells did not display LTC_4_ synthesizing activity (**Fig 4C** top image; LTC_4_^-^ mononuclear cells indicated by asterisks). In fact, DP2 receptors appear to regulate neither migration nor activation of peritoneal mononuclear cells in this mice model of *S. mansoni* infection, since CAY10471 treatment, besides LTC_4_ synthesis, also did not alter *S. mansoni* infection-associated (i) numbers of peritoneal mononuclear cells (mixture of mast cells, monocytes/macrophages and lymphocytes) found 8 wpi (**S2A Fig**); (ii) the scarce cytoplasmic lipid bodies found in peritoneal mononuclear cells (**S2B Fig**) compared to elevated content within eosinophils (**Fig 4B**); and (iii) lack of intracellular immunolabelling for newly synthesized LTC_4_ (**Fig 4C**). Therefore, eosinophils seem to represent the cell source of PGD_2_/DP2 receptor-driven cysLTs produced in the peritoneal compartment of *S. mansoni* infected animals.

Moving from the peritoneal compartment to the granulomatous liver of *S. mansoni*-infected mice, CAY10471 treatment also promoted the reduction of total hepatic cysLTs (LTC_4_/D_4_/E_4_) content within the *S. mansoni* infection-induced eosinophilic granuloma-enriched liver (**Fig 5A**). CAY10471 also produced an even more robust inhibition than those observed with HQL-79 treatment on the magnitude of eosinophilia within hepatic granulomas found 8 wpi (**Fig 5A**). A positive correlation (r^2^ = 0.7253; *p* ≤ 0.0018) derived from this data indicates a product/producer relationship between cysLTs and granulomatous eosinophils under CAY10471 treatment (**Fig 5B**). Taking together with HQL-79 data, we speculate that an *S. mansoni* infection-induced DP2 activation by PGD_2_ stimulates *in situ* LTC_4_ synthesis/release by eosinophils infiltrating hepatic granulomas, therefore bringing about the increased liver content of cysLTs.

**Fig 5.**
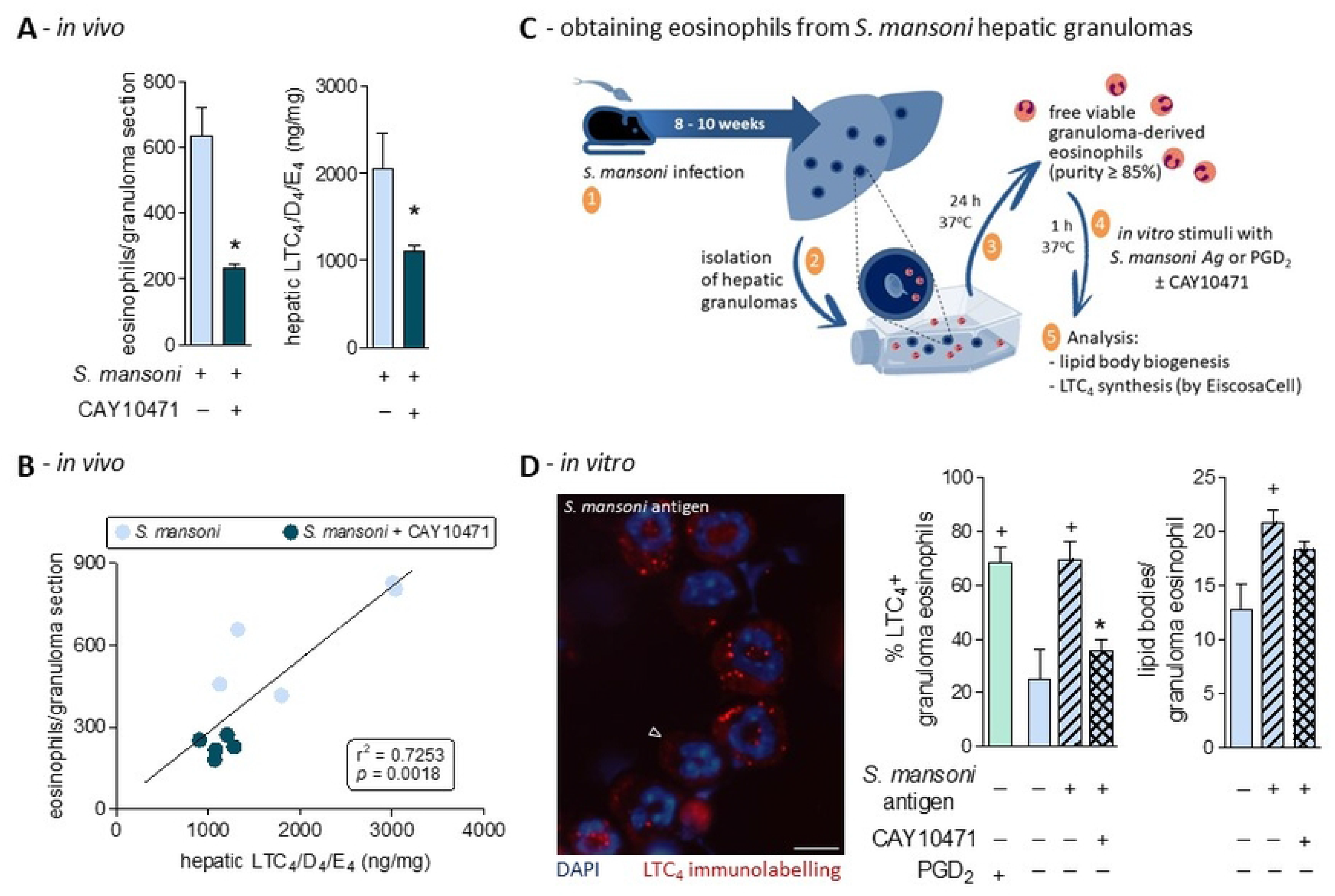
DP2 receptor activation triggers *S. mansoni* granuloma eosinophil response: PGD_2_-stimulated eosinophils as the cellular source of DP2-driven *de novo* synthesized LTC_4._ **A** shows enumeration of granuloma-infiltrating eosinophils as well as the release of cysLTs (LTC_4_/D_4_/E_4_) detected by specific ELISA kits in liver homogenates. Values are expressed as the mean ± SEM from at least 5 animals *per* group (experiment was repeated at least once). ^+^*p* < 0.05 compared to non-infected control group. \**p* < 0.05 compared to infected non-treated group. **B** displays linear regression curve illustrating the relationship between cysLTS levels and granulomatous eosinophilic reaction found in hepatic tissues. R-squared (r^2^) and *p* values for the linear regression are shown. Panel **C** schematizes protocol for the isolation of eosinophils from schistosomal hepatic granulomas (a cellular suspension displaying purity of about 85 to 90%). Image in **D** shows a representative image of intracellularly immunolabeled LTC_4_ (in red; for EicosaCell preparation *vide* Methods) within granuloma-isolated eosinophils stimulated *in vitro* for 1 h with *S. mansoni* antigen (0.5 µg/mL). Cellular nuclei are labeled with DAPI (in blue); bar = 10 µm; arrowhead indicates an LTC_4_^-^ eosinophil. Graphs in **D** show the percentage of *in vitro* stimulated eosinophils (as indicated) exhibiting cytoplasmic immunostaining for LTC_4_ (LTC_4_^+^ eosinophils) as well as the cytoplasmic numbers of eosinophil lipid body organelles (evaluated in osmium-stained cells). Values are expressed as the mean ± SEM from 5 preparations of granuloma-isolated eosinophils. ^+^*p* < 0.05 compared to non-stimulated eosinophils. \**p* < 0.05 compared to *S. mansoni* antigen-stimulated eosinophils

Despite the correlation between eosinophil infiltration and cysLT content in the liver, a demonstration that granulomatous eosinophils indeed generate LTC_4_ was still to be determined. To investigate whether the eosinophils present in the fibrotic hepatic granulomas were LTC_4_ synthesizing cells accountable for the increased cysLTs levels found in schistosomal livers, we isolated eosinophils from the *S. mansoni* hepatic granulomas, stimulated them *in vitro* with *S. mansoni* antigen or PGD_2_ for 1 h (with or without CAY10471) and then analyzed lipid body biogenesis as well as intracellular LTC_4_ synthesis (**Fig 5C**). In this *in vitro* experimental setting, direct stimulations with PGD_2_ (25 nM) triggered rapid (1 h) intracellular LTC_4_ synthesis within about 70% of granuloma-isolated eosinophils (**Fig 5D**). So, hepatic granulomatous eosinophils under PGD_2_ stimulation are indeed capable of LTC_4_ synthesis. Moreover, *in vitro* co-incubation of granuloma-isolated eosinophils with CAY10471 blocked *S. mansoni* antigen ability to induce LTC_4_ synthesis (**Fig 5D**). So, at least *in vitro*, an autocrine/paracrine phenomenon takes place to regulate LTC_4_ synthesis by *S. mansoni* granuloma-derived eosinophils. However, whether such a DP2-dependent autocrine phenomenon controls eosinophil-driven LTC_4_ synthesis *in vivo* depends on a prospective characterization of PGD_2_ cell source in a hepatic granulomatous environment.

For either human blood or in murine pleural fluid cells, eosinophil lipid bodies are the primary subcellular site of PGD_2_-induced LTC_4_ synthesis [12,11,64]. As shown in **Fig 5D**, immunolabelling of newly synthesized LTC_4_ was in a punctate pattern, with cytoplasmic distribution apart from the perinuclear membrane and fully consistent in size and numbers with eosinophil lipid bodies. As confirmed here (**Fig 5D**, right graph), *in vitro* co-incubation with CAY10471 does not modify *S. mansoni* antigen-triggered lipid body biogenesis, a PGD_2_-induced intracellular event downstream to DP1 receptor activation, rather than DP2 [15]. Besides eosinophils, various cell types comprising *S. mansoni*-driven hepatic granuloma can synthesize LTC_4_, including hepatic stellate myofibroblasts. However, in response to PGD_2_, eosinophils are the only characterized cells that synthesize LTC_4_ within cytoplasmic lipid bodies [10]. The remaining question of whether additional cells, besides eosinophils, are contributing to the global PGD_2_-induced cysLTs production during *S. mansoni* infection still needs to be addressed.

The two direct attempts to define eosinophil function in schistosomiasis employing mouse models of eosinophil lineage deficiency, including the studies by Swartz et al. (2006) and de Oliveira et al. (2022), did not uphold the original assumption of eosinophil effector cytotoxic function towards parasites. Instead, these pivotal studies have unveiled a more subtle immunomodulatory role for eosinophils in *S. mansoni*-evoked granulomatous pathogenesis [48,49]. Particularly regarding *S. mansoni* infection-triggered hepatic fibrosis within eosinophilic granulomas, de Oliveira and coauthors (2022) revealed that in the absence of eosinophils, reductions in TGF-β and IL-13 levels were accompanied by attenuation of hepatic fibrosis [48], thus indicating a pro-fibrogenic role for eosinophils in the disease. These findings are in clear contrast with the inverse relationship between *S. mansoni* infection-related eosinophilia and fibrogenesis observed here, whereby decreasing PGD_2_-driven eosinophilia (**Fig 3** and **4**), HQL-79 and CAY-10471 did increase the fibrogenic cytokines as well as hepatic fibrosis (**Fig 2**). It seems that targeting specifically PGD_2_-regulated eosinophil functions after initial evets of infection, rather than the entire eosinophil population from the beginning, appears to promote a distinct pattern of response in schistosomiasis. It is also noteworthy that by employing virtually the same transgenic model of eosinophil ablation, the study by Swartz and colleagues (2006) uncovered an atypical but prominent mast cell infiltration within eosinophil-free granulomas of eosinophil deficient mice [49]. As neither eosinophil ablation study evaluated PGD_2_ component of schistosomal fibrogenic response [49,48], one may speculate that such mast cell phenomenon may represent a counterbalancing response with anti-fibrogenic outcomes, perhaps mediated by PGD_2_. Mast cells are skilled producers of PGD_2_ [65], which here was identified as an endogenous down-regulatory signal of *S. mansoni*-triggered hepatic granulomatous fibrosis. Furthermore, mast cells are also important cellular sources of LTC_4_ [65]. So, one can speculate that functioning as eosinophil substitutes in the hepatic granulomatous tissue of S*. mansoni*-infected eosinophil deficient mice, mast cells may secrete PGD_2_ and/or LTC_4_ which appears to be related to down-regulatory effects in schistosomal hepatic fibrosis (**Fig 4C**; [56])

## Conclusions

The detrimental effect of both HQL-79 and CAY10471 on *S. mansoni* infection-driven hepatic granulomatous inflammation uncovered here has disclosed an unanticipated endogenous protective mechanism of the hepatic tissue – the *in vivo* anti-fibrogenic activity of hPGDS-derived PGD_2_ via DP2 activation. In addition, our pharmacological approaches have also unveiled a potential novel PGD_2_-driven regulatory mechanism: DP2 receptors-elicited eosinophils as a major cellular source of hepatic LTC_4_ in *S. mansoni* infection. Future investigations should focus to define eosinophil-derived cysLTs role in *S. mansoni* infection-elicited hepatic fibrosis by pharmacological selective strategies, rather than genetic approaches that affect initial cercaria/schistosomula-driven initial events. For instance, these studies may show cysLTs as important immunoregulators of IL-4-mediated type 2 immune response of schistosomiasis, since LTC_4_ is known to function as an intracrine signal capable of triggering IL-4 secretion from eosinophils [66,67].

A main driver for the current study was the *in vitro* data on PGD_2_ role in hepatic tissue-related fibrosis suggesting that PGD_2_ displays deleterious functions in the context of *S. mansoni* infection [21,8]; and more importantly, the resulting recommendations for PGD_2_-targeting therapeutic maneuvers in schistosomiasis [33–35]. Because presently there are other clinically approved uses for DP2 receptor antagonism in type 2 immune response-biased pathologies [i.e., ramatroban (Baynas®) in allergic rhinitis], our current study compels that drug repurposing (i) should not arise directly from the *in vitro* studies, and most importantly (ii) should be avoided in this case. Concurring, the mechanism involved in DP2 receptor anti-fibrogenic function has recently emerged as a far more complex activity. For instance, it has been shown that the DP2 molecule appears to traffic back and anchor to endoplasmic reticulum membranes where, in a fashion non-related to PGD_2_, DP2 evokes degradation of collagen mRNA and decreases intracellular collagen biosynthesis [68]. Also, data derived from genetically-deficient animals should be carefully analyzed before being incorporated into treatment strategies as they may not reflect the proper window of opportunity for the infected patient at risk of developing complications due to hepatic fibrosis. Indeed, based on our current *in vivo* findings, promoting pharmacological DP2 selective agonism (not antagonism) may be the right track to achieve anti-fibrogenic effects.

## Materials and methods

### Animals and ethics statement

Male C57BL/6 mice weighing 20 to 25 g were obtained from either CCS/UFRJ or FIOCRUZ breeding centers, raised, and maintained under the same housing conditions. Animals were housed in a temperature-controlled room, under a 12∶12 h light cycle with access to filtered water and chow *ad libitum*. All animal care and experimental protocols were conducted following the guidelines of the Brazilian Council for Care and Use of Experimentation Animals (CONCEA) in accordance with Brazilian Federal Law number 11.794/2008, which regulates the scientific use of animals (license numbers CEUA115/14 and CEUA139/21 by the Committee for Ethics in Animal Experimentation of Federal University of Rio de Janeiro).

### *S. mansoni* infection and treatments

Mice were infected by active percutaneous penetration of *S. mansoni* infective stage (BH strain; Institute Oswaldo Cruz, FIOCRUZ, RJ), by exposition of washed mouse paws to 60 alive cercariae for 40 min. Uninfected age-matched mice were used as controls. At 3.5 weeks post infection (wpi), osmotic pumps (Alzet® pump; flow rate of 0.11 μL/hour - as stated by the manufacturer) – containing 100 µL of either HQL-79 (1 mg/mL) or CAY10471 (670 µg/mL) solutions producing a cumulative dose of 2.6 or 1.7 µg/day, respectively – were implanted subcutaneously in infected and non-infected mice. Both HQL-79 and CAY10471 were diluted with 0.1 % DMSO in sterile saline. Osmotic pumps containing vehicle solution implanted subcutaneously did not cause changes in mice survival rate, hepatic disfunction, or inflammatory alterations within implant sites, peripheral blood or peritoneal cavities of either non-infected or infected mice (data not shown). All animals were euthanized under anesthesia after 8 wpi and during this time, they were maintained under same housing conditions described before.

### Analysis of parasitological parameters

To ascertain the establishment of the infection and characterize oviposition onset, the presence of *S. mansoni* eggs in feces and hepatic tissue were determined. *S. mansoni* eggs were identified by morphology (containing sharp lateral spines) and enumerated by copro-parasitological thick-smear Kato-Katz method (Helm-Test Biomanguinhos; FIOCRUZ), while the presence of *S. mansoni* eggs found trapped in hepatic granulomas was analyzed in liver histological preparations (as described below).

### Histopathological analysis

Liver samples were washed in saline, fixed in 10% buffered formalin, dehydrated in alcohol, and embedded in paraffin. Sections (5 μm of thickness) were stained with (i) hematoxylin-eosin for enumeration and area analysis of granulomas with egg entrapped; (ii) modified PicroSirius technique for collagen fibers deposition detection; or (iii) Sirius Red for granuloma-infiltrating eosinophils quantification. Acquisition of digital photographs and quantitative analysis of hepatic tissue sections were carried out using a slider scanner (Pannoramic Midi 3DHistec or Motic Easy Scan) and a computer-assisted image analyzer (Slide Viewer 2.4 or Qu Path 0.4.3). All evaluations were performed by two different blinded observers.

Under light microscopy, the areas of hepatic granulomas (20 granulomas *per* animal) were determined in digital images (acquired at 20x magnification) of histological sections, containing central eggs, randomly chosen, using the measurement tool of the image analyzer. Egg-encasing granulomas counting was carried out at a low power (20x) magnification under light microscopy. The mean number of egg-encasing granulomas *per* field was calculated for each infected mouse. By confocal microscopy, total fluorescent intensity within digital images of individual granuloma areas (10 granulomas *per* animal) was quantified using Image J software. All eosinophils identified in individual granuloma areas were counted in digital images acquired under 80x magnification light microscopy. Ten random granulomas *per* animal were analyzed and results were expressed as eosinophils/granuloma.

### Peripheral blood, peritoneal, and bone marrow eosinophilia

Blood eosinophilia was analyzed by light microscopy of blood smears stained with Panoptic kit. To evaluate peritoneal and bone marrow eosinophilia, cells from peritoneal cavities and bone marrows of removed femurs were harvested with RPMI 1640 medium (Sigma) and cytospun towards slides. Total leukocyte enumeration was performed in Neubauer chambers and differential eosinophil count in Panoptic kit-stained slides under light microscopy.

### Eosinophils isolation from *S. mansoni*-driven hepatic granuloma

Mice livers recovered from 8 to 10 wpi with *S. mansoni* were homogenized in RPMI 1640 medium. Intact hepatic granulomas were allowed to sediment, washed 3 times with RPMI, and then incubated overnight (37°C; 5% CO_2_). In culture bottles, granuloma-released eosinophils are non-adherent cells in suspension, while other cells are found still within granulomas or attached to the plastic. Recovered cells were analyzed under light microscopy after Panoptic kit staining to determine the percentage of eosinophils in suspension. The eosinophil fraction used in *in vitro* experiments was composed by 80 to 90% purified eosinophils.

Isolated eosinophils (3 x 10^6^ cells/mL) were co-incubated or not with CAY10471 (200 nM; Cayman Chemicals) and then stimulated for 1 h (37 °C; 5% CO_2_) with *S. mansoni* antigen (0.5 µg/mL; Cusabio) or PGD_2_ (25 nM; Cayman Chemicals). After incubation, eosinophil samples were promptly fixed with PFO for osmium staining or placed in EDAC solution for EicosaCell assay (see below).

### Evaluation of hepatic collagen synthesis

As an indirect quantitative assay determining amounts of collagen molecules, hepatic fibrosis was also evaluated by measuring hydroxyproline levels in liver homogenates.

Hydroxyproline content was determined in dehydrated/hydrolyzed liver fragments by a colorimetric method in chloramine-T buffer (Sigma, USA) with Ehrlich’s reagent (Sigma)/perchloric acid (Merck) and absorbance was measured at 557 nm. Results were expressed as ng of hydroxyproline *per* mg of hepatic tissue.

### Analysis of lipid mediators production

Amounts of eicosanoids PGD_2_, PGE_2_, and cysteinyl leukotrienes (LTC_4_/D_4_/E_4_) found in peritoneal fluid and/or homogenized liver fragments were measured by specific commercial EIA kits, according to the manufacturer’s instructions (Cayman Chemicals).

For cysLTs production, intracellular LTC_4_ synthesis was alternatively analyzed by the EicosaCell methodology [69] – an immuno-assay that immobilizes the newly formed eicosanoid within active synthesizing cells. Briefly, peritoneal cells or *in vitro*-stimulated granuloma-derived eosinophils (cell suspensions in RPMI) were immediately mixed with an equal volume of a 0.2% solution of 1-ethyl-3-(3-dimethylamino-propyl) carbodiimide (EDAC; in PBS), used to crosslink eicosanoid carboxyl groups to amines in proteins. After a 10 min incubation at room temperature with EDAC, eosinophils were washed with PBS, cytospun onto glass slides, fixed with PFO (2%), incubated with PBS with 1% BSA for 15 min, and then incubated with a rabbit anti-LTC_4_ antibody (Cayman Chemicals) overnight. The cells were washed with PBS containing 1% BSA (3 times 10 min) and then incubated with Alexa594 donkey anti-rabbit secondary IgG (Jackson) for 1 h. Nuclear visualization by DAPI staining was employed to distinguish polymorphonuclear eosinophils from mononuclear cells. EicosaCell images were obtained using an Olympus BX51 fluorescence microscope, equipped with a 100X objective in conjunction with LAS-AF 2.2.0 Software.

### Cytokines measurements

Amounts of TGF-β, IL-5, and IL-13 in liver fragments or in serum samples were measured by commercial ELISA kits, according to the manufacturer’s instructions (R&D Systems and/or Peprotech).

### Lipid Body Staining and Enumeration

For lipid body counting, cells in cytospin slides were fixed in 3.7% paraformaldehyde and stained with 1.5% OsO4 in 0.1 M cacodylate buffer, as previously described [64]. By bright field microscopy, fifty consecutively eosinophils/slide were evaluated in a blinded fashion by more than one observer.

### Statistical analysis

Results are expressed as means ± SEM (standard error of the mean) and were analyzed by one-way ANOVA, followed by Student-Newman-Keuls test or by Student’s *t* test, using Prism software (GraphPad Software, Inc., San Diego, CA, USA). Differences were considered significant when *p* < 0.05. Two independent experiments were performed for each PGD_2_-targeting treatment studied: HQL-79 and CAY10471.

---

**S1 Fig.**
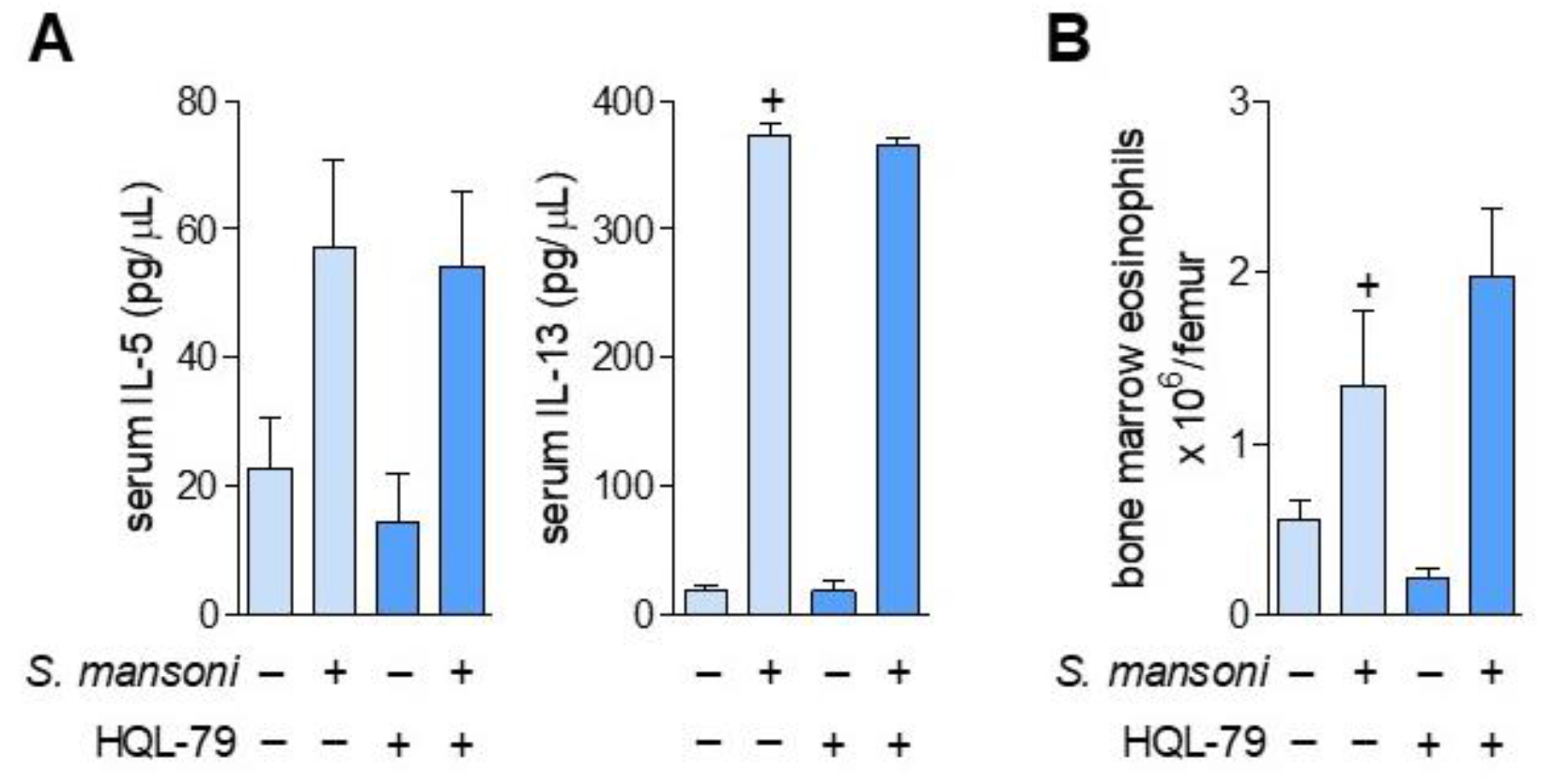
Inhibition of PGD_2_ synthesis by HQL-79 treatment did not affect *S. mansoni* infection-induced immune polarization to a type 2-biased response in mice. **A** shows serum levels of IL-5 and IL-13. B shows eosinophil numbers found at bone marrow. Values are expressed as the mean ± SEM from at least 5 animals *per* group (experiment was repeated at least once). ^+^*p* < 0.05 compared to non-infected control group.

**S2 Fig.**
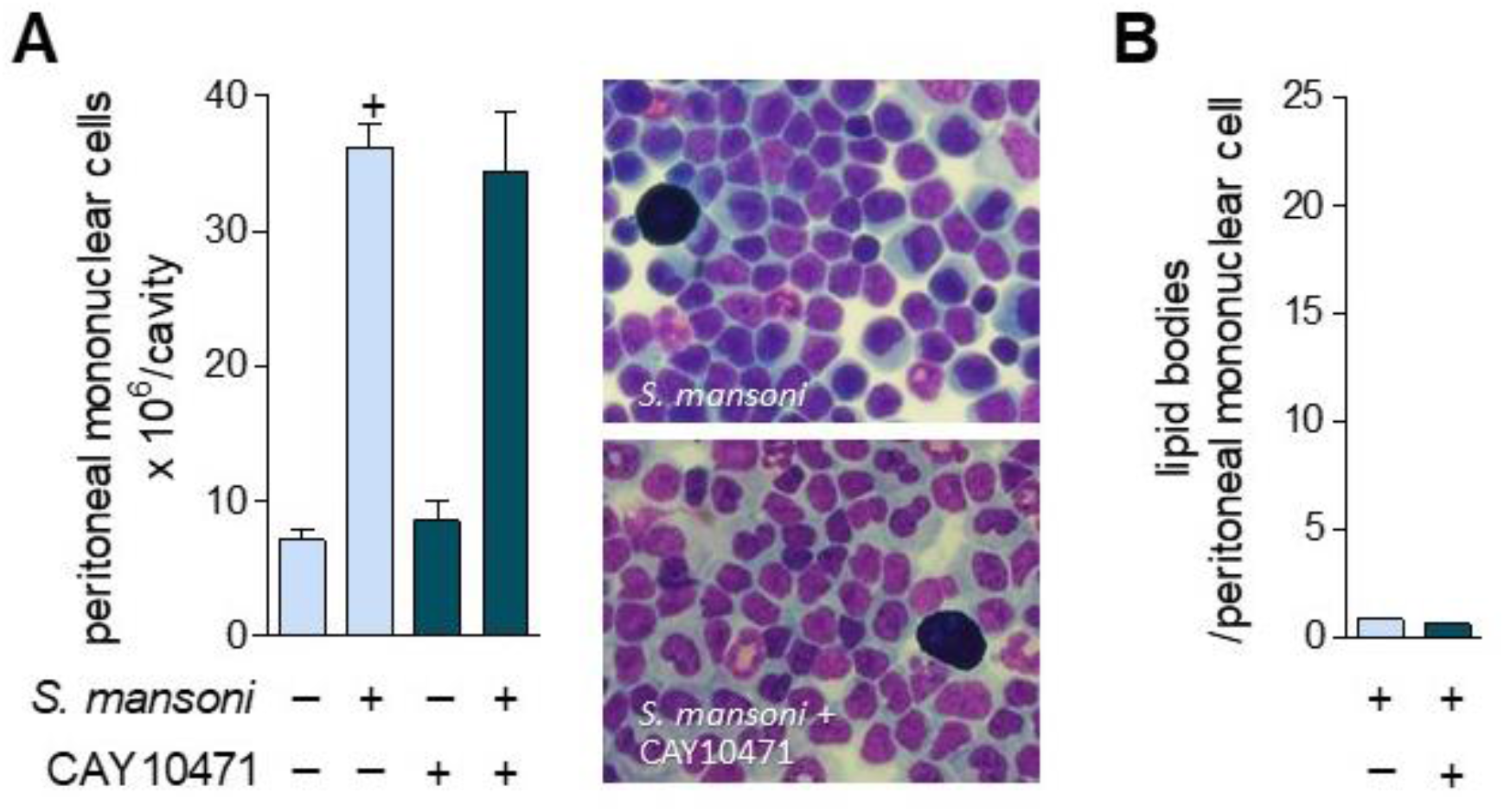
Antagonism of DP2 receptor by CAY10471 treatment did not affect *S. mansoni* infection-elicited population of peritoneal mononuclear cells. **A** shows numbers of mononuclear cells found at peritoneal space as well as representative images of peritoneal cells 8 wpi of *S. mansoni*-infected mice (top image) and in mice treated with CAY10471 and *S. mansoni* infected (bottom image). **B** shows numbers of cytoplasmic lipid body organelles found in peritoneal mononuclear cells. Values are expressed as the mean ± SEM from at least 5 animals *per* group (experiment was repeated at least once). ^+^*p* < 0.05 compared to non-infected control group.

## Notes

### Competing Interest Statement

The authors have declared no competing interest.

## References

1. Zwang J, Olliaro P. Efficacy and safety of praziquantel 40 mg/kg in preschool-aged and school-aged children: a meta-analysis. Parasit Vectors. 2017;10: 47. doi:10.1186/s13071-016-1958-7

2. Zaparina O, Rakhmetova AS, Kolosova NG, Cheng G, Mordvinov VA, Pakharukova MY. Antioxidants resveratrol and SkQ1 attenuate praziquantel adverse effects on the liver in Opisthorchis felineus infected hamsters. Acta Trop. 2021;220: 105954. doi:10.1016/j.actatropica.2021.105954

3. Niu X, Hu T, Hong Y, Li X, Shen Y. The Role of Praziquantel in the Prevention and Treatment of Fibrosis Associated with Schistosomiasis: A Review. J Trop Med. 2022;2022: 1413711. doi:10.1155/2022/1413711

4. Fate tracing reveals hepatic stellate cells as dominant contributors to liver fibrosis independent of its aetiology - PubMed. [cited 6 Sep 2023]. Available: https://pubmed.ncbi.nlm.nih.gov/24264436/

5. Anthony B, Mathieson W, de Castro-Borges W, Allen J. Schistosoma mansoni: egg-induced downregulation of hepatic stellate cell activation and fibrogenesis. Exp Parasitol. 2010;124: 409–420. doi:10.1016/j.exppara.2009.12.009

6. Chiaramonte MG, Cheever AW, Malley JD, Donaldson DD, Wynn TA. Studies of murine schistosomiasis reveal interleukin-13 blockade as a treatment for established and progressive liver fibrosis. Hepatology. 2001;34: 273–282. doi:10.1053/jhep.2001.26376

7. Chiaramonte MG, Donaldson DD, Cheever AW, Wynn TA. An IL-13 inhibitor blocks the development of hepatic fibrosis during a T-helper type 2-dominated inflammatory response. J Clin Invest. 1999;104: 777–785. doi:10.1172/JCI7325

8. Paiva LA, Coelho KA, Luna-Gomes T, El-Cheikh MC, Borojevic R, Perez SA, et al. Schistosome infection-derived Hepatic Stellate Cells are cellular source of prostaglandin D₂: role in TGF-β-stimulated VEGF production. Prostaglandins Leukot Essent Fatty Acids. 2015;95: 57–62. doi:10.1016/j.plefa.2015.01.004

9. Rittchen S, Heinemann A. Therapeutic Potential of Hematopoietic Prostaglandin D2 Synthase in Allergic Inflammation. Cells. 2019;8. doi:10.3390/cells8060619

10. Luna-Gomes T, Magalhães KG, Mesquita-Santos FP, Bakker-Abreu I, Samico RF, Molinaro R, et al. Eosinophils as a novel cell source of prostaglandin D2: autocrine role in allergic inflammation. J Immunol. 2011;187: 6518–6526. doi:10.4049/jimmunol.1101806

11. Amorim NRT, Luna-Gomes T, Gama-Almeida M, Souza-Almeida G, Canetti C, Diaz BL, et al. Leptin Elicits LTC4 Synthesis by Eosinophils Mediated by Sequential Two-Step Autocrine Activation of CCR3 and PGD2 Receptors. Front Immunol. 2018;9. doi:10.3389/fimmu.2018.02139

12. Luna-Gomes T, Bozza PT, Bandeira-Melo C. Eosinophil recruitment and activation: the role of lipid mediators. Front Pharmacol. 2013;4: 27. doi:10.3389/fphar.2013.00027

13. Jandl K, Heinemann A. The therapeutic potential of CRTH2/DP2 beyond allergy and asthma. Prostaglandins Other Lipid Mediat. 2017;133: 42–48. doi:10.1016/j.prostaglandins.2017.08.006

14. Monneret G, Gravel S, Diamond M, Rokach J, Powell WS. Prostaglandin D2 is a potent chemoattractant for human eosinophils that acts via a novel DP receptor. Blood. 2001;98: 1942–1948. doi:10.1182/blood.v98.6.1942

15. Mesquita-Santos FP, Bakker-Abreu I, Luna-Gomes T, Bozza PT, Diaz BL, Bandeira-Melo C. Co-operative signalling through DP(1) and DP(2) prostanoid receptors is required to enhance leukotriene C(4) synthesis induced by prostaglandin D(2) in eosinophils. Br J Pharmacol. 2011;162: 1674–1685. doi:10.1111/j.1476-5381.2010.01086.x

16. Peinhaupt M, Roula D, Theiler A, Sedej M, Schicho R, Marsche G, et al. DP1 receptor signaling prevents the onset of intrinsic apoptosis in eosinophils and functions as a transcriptional modulator. J Leukoc Biol. 2018;104: 159–171. doi:10.1002/JLB.3MA1017-404R

17. Mesquita-Santos FP, Vieira-de-Abreu A, Calheiros AS, Figueiredo IH, Castro-Faria-Neto HC, Weller PF, et al. Cutting edge: prostaglandin D2 enhances leukotriene C4 synthesis by eosinophils during allergic inflammation: synergistic in vivo role of endogenous eotaxin. J Immunol. 2006;176: 1326–1330. doi:10.4049/jimmunol.176.3.1326

18. Fusco AC, Salafsky B, Kevin MB. Schistosoma mansoni: eicosanoid production by cercariae. Exp Parasitol. 1985;59: 44–50. doi:10.1016/0014-4894(85)90055-4

19. Hervé M, Angeli V, Pinzar E, Wintjens R, Faveeuw C, Narumiya S, et al. Pivotal roles of the parasite PGD2 synthase and of the host D prostanoid receptor 1 in schistosome immune evasion. Eur J Immunol. 2003;33: 2764–2772. doi:10.1002/eji.200324143

20. Johnson KA, Angelucci F, Bellelli A, Hervé M, Fontaine J, Tsernoglou D, et al. Crystal structure of the 28 kDa glutathione S-transferase from Schistosoma haematobium. Biochemistry. 2003;42: 10084–10094. doi:10.1021/bi034449r

21. Magalhães KG, Luna-Gomes T, Mesquita-Santos F, Corrêa R, Assunção LS, Atella GC, et al. Schistosomal Lipids Activate Human Eosinophils via Toll-Like Receptor 2 and PGD2 Receptors: 15-LO Role in Cytokine Secretion. Front Immunol. 2018;9: 3161. doi:10.3389/fimmu.2018.03161

22. Angeli V, Faveeuw C, Roye O, Fontaine J, Teissier E, Capron A, et al. Role of the parasite-derived prostaglandin D2 in the inhibition of epidermal Langerhans cell migration during schistosomiasis infection. J Exp Med. 2001;193: 1135–1147. doi:10.1084/jem.193.10.1135

23. Zhang A, Dong Z, Yang T. Prostaglandin D2 inhibits TGF-beta1-induced epithelial-to-mesenchymal transition in MDCK cells. Am J Physiol Renal Physiol. 2006;291: F1332–1342. doi:10.1152/ajprenal.00131.2006

24. Takahashi N, Kikuchi H, Usui A, Furusho T, Fujimaru T, Fujiki T, et al. Deletion of Alox15 improves kidney dysfunction and inhibits fibrosis by increased PGD2 in the kidney. Clin Exp Nephrol. 2021;25: 445–455. doi:10.1007/s10157-021-02021-y

25. Ito H, Yan X, Nagata N, Aritake K, Katsumata Y, Matsuhashi T, et al. PGD2-CRTH2 pathway promotes tubulointerstitial fibrosis. J Am Soc Nephrol. 2012;23: 1797–1809. doi:10.1681/ASN.2012020126

26. Ayabe S, Kida T, Hori M, Ozaki H, Murata T. Prostaglandin D2 inhibits collagen secretion from lung fibroblasts by activating the DP receptor. J Pharmacol Sci. 2013;121: 312–317. doi:10.1254/jphs.12275fp

27. Kida T, Ayabe S, Omori K, Nakamura T, Maehara T, Aritake K, et al. Prostaglandin D2 Attenuates Bleomycin-Induced Lung Inflammation and Pulmonary Fibrosis. PLoS One. 2016;11: e0167729. doi:10.1371/journal.pone.0167729

28. Ueda S, Fukunaga K, Takihara T, Shiraishi Y, Oguma T, Shiomi T, et al. Deficiency of CRTH2, a Prostaglandin D2 Receptor, Aggravates Bleomycin-induced Pulmonary Inflammation and Fibrosis. Am J Respir Cell Mol Biol. 2019;60: 289–298. doi:10.1165/rcmb.2017-0397OC

29. Lacy SH, Epa AP, Pollock SJ, Woeller CF, Thatcher TH, Phipps RP, et al. Activated human T lymphocytes inhibit TGFβ-induced fibroblast to myofibroblast differentiation via prostaglandins D2 and E2. Am J Physiol Lung Cell Mol Physiol. 2018;314: L569–L582. doi:10.1152/ajplung.00565.2016

30. Brightling CE, Brusselle G, Altman P. The impact of the prostaglandin D2 receptor 2 and its downstream effects on the pathophysiology of asthma. Allergy. 2020;75: 761–768. doi:10.1111/all.14001

31. Pelaia C, Crimi C, Vatrella A, Busceti MT, Gaudio A, Garofalo E, et al. New treatments for asthma: From the pathogenic role of prostaglandin D2 to the therapeutic effects of fevipiprant. Pharmacol Res. 2020;155: 104490. doi:10.1016/j.phrs.2019.104490

32. Paiva LA, Maya-Monteiro CM, Bandeira-Melo C, Silva PMR, El-Cheikh MC, Teodoro AJ, et al. Interplay of cysteinyl leukotrienes and TGF-β in the activation of hepatic stellate cells from Schistosoma mansoni granulomas. Biochim Biophys Acta. 2010;1801: 1341–1348. doi:10.1016/j.bbalip.2010.08.014

33. Coakley G, Wright MD, Borger JG. Schistosoma mansoni-Derived Lipids in Extracellular Vesicles: Potential Agonists for Eosinophillic Tissue Repair. Front Immunol. 2019;10: 1010. doi:10.3389/fimmu.2019.01010

34. Acharya S, Da’dara AA, Skelly PJ. Schistosome immunomodulators. PLoS Pathog. 2021;17: e1010064. doi:10.1371/journal.ppat.1010064

35. Fan S, Chen W, Chen L, Li L. CRTH2: a potential target for the treatment of organ fibrosis. Acta Biochim Biophys Sin (Shanghai). 2022;54: 590–592. doi:10.3724/abbs.2022025

36. Murata T, Aritake K, Tsubosaka Y, Maruyama T, Nakagawa T, Hori M, et al. Anti-inflammatory role of PGD2 in acute lung inflammation and therapeutic application of its signal enhancement. Proc Natl Acad Sci U S A. 2013;110: 5205–5210. doi:10.1073/pnas.1218091110

37. Kobayashi K, Tsubosaka Y, Hori M, Narumiya S, Ozaki H, Murata T. Prostaglandin D2-DP signaling promotes endothelial barrier function via the cAMP/PKA/Tiam1/Rac1 pathway. Arterioscler Thromb Vasc Biol. 2013;33: 565–571. doi:10.1161/ATVBAHA.112.300993

38. Muniz VS, Baptista-Dos-Reis R, Benjamim CF, Mata-Santos HA, Pyrrho AS, Strauch MA, et al. Purinergic P2Y12 Receptor Activation in Eosinophils and the Schistosomal Host Response. PLoS One. 2015;10: e0139805. doi:10.1371/journal.pone.0139805

39. Pearce EJ, MacDonald AS. The immunobiology of schistosomiasis. Nat Rev Immunol. 2002;2: 499–511. doi:10.1038/nri843

40. Luna-Gomes T, Bozza PT, Bandeira-Melo C. Eosinophil recruitment and activation: the role of lipid mediators. Front Pharmacol. 2013;4: 27. doi:10.3389/fphar.2013.00027

41. Jandl K, Stacher E, Bálint Z, Sturm EM, Maric J, Peinhaupt M, et al. Activated prostaglandin D2 receptors on macrophages enhance neutrophil recruitment into the lung. J Allergy Clin Immunol. 2016;137: 833–843. doi:10.1016/j.jaci.2015.11.012

42. Wu D, Molofsky AB, Liang H-E, Ricardo-Gonzalez RR, Jouihan HA, Bando JK, et al. Eosinophils sustain adipose alternatively activated macrophages associated with glucose homeostasis. Science. 2011;332: 243–247. doi:10.1126/science.1201475

43. Mederacke I, Hsu CC, Troeger JS, Huebener P, Mu X, Dapito DH, et al. Fate tracing reveals hepatic stellate cells as dominant contributors to liver fibrosis independent of its aetiology. Nat Commun. 2013;4: 2823. doi:10.1038/ncomms3823

44. Anthony B, Allen JT, Li YS, McManus DP. Hepatic stellate cells and parasite-induced liver fibrosis. Parasit Vectors. 2010;3: 60. doi:10.1186/1756-3305-3-60

45. Goh YPS, Henderson NC, Heredia JE, Red Eagle A, Odegaard JI, Lehwald N, et al. Eosinophils secrete IL-4 to facilitate liver regeneration. Proc Natl Acad Sci USA. 2013;110: 9914–9919. doi:10.1073/pnas.1304046110

46. Xu L, Yang Y, Wen Y, Jeong J-M, Emontzpohl C, Atkins CL, et al. Hepatic recruitment of eosinophils and their protective function during acute liver injury. J Hepatol. 2022;77: 344–352. doi:10.1016/j.jhep.2022.02.024

47. Weller PF, Spencer LA. Functions of tissue-resident eosinophils. Nat Rev Immunol. 2017;17: 746–760. doi:10.1038/nri.2017.95

48. de Oliveira VG, Rodrigues VF, Moreira JMP, Rodrigues JL, Maggi L, Resende SD, et al. Eosinophils participate in modulation of liver immune response and tissue damage induced by Schistosoma mansoni infection in mice. Cytokine. 2022;149: 155701. doi:10.1016/j.cyto.2021.155701

49. Swartz JM, Dyer KD, Cheever AW, Ramalingam T, Pesnicak L, Domachowske JB, et al. Schistosoma mansoni infection in eosinophil lineage-ablated mice. Blood. 2006;108: 2420–2427. doi:10.1182/blood-2006-04-015933

50. Malta KK, Palazzi C, Neves VH, Aguiar Y, Silva TP, Melo RCN. Schistosomiasis Mansoni-Recruited Eosinophils: An Overview in the Granuloma Context. Microorganisms. 2022;10: 2022. doi:10.3390/microorganisms10102022

51. Hirai H, Tanaka K, Yoshie O, Ogawa K, Kenmotsu K, Takamori Y, et al. Prostaglandin D2 selectively induces chemotaxis in T helper type 2 cells, eosinophils, and basophils via seven-transmembrane receptor CRTH2. J Exp Med. 2001;193: 255–261. doi:10.1084/jem.193.2.255

52. Spencer LA, Szela CT, Perez SAC, Kirchhoffer CL, Neves JS, Radke AL, et al. Human eosinophils constitutively express multiple Th1, Th2, and immunoregulatory cytokines that are secreted rapidly and differentially. J Leukoc Biol. 2009;85: 117–123. doi:10.1189/jlb.0108058

53. Al-Azzam N, Elsalem L. Leukotriene D4 role in allergic asthma pathogenesis from cellular and therapeutic perspectives. Life Sci. 2020;260: 118452. doi:10.1016/j.lfs.2020.118452

54. Kanaoka Y, Boyce JA. Cysteinyl leukotrienes and their receptors: cellular distribution and function in immune and inflammatory responses. J Immunol. 2004;173: 1503–1510. doi:10.4049/jimmunol.173.3.1503

55. Peters-Golden M. Expanding roles for leukotrienes in airway inflammation. Curr Allergy Asthma Rep. 2008;8: 367–373. doi:10.1007/s11882-008-0057-z

56. Toffoli da Silva G, Espíndola MS, Fontanari C, Rosada RS, Faccioli LH, Ramos SG, et al. 5-lipoxygenase pathway is essential for the control of granuloma extension induced by Schistosoma mansoni eggs in lung. Exp Parasitol. 2016;167: 124–129. doi:10.1016/j.exppara.2016.06.001

57. Machado ER, Ueta MT, Lourenço EV, Anibal FF, Sorgi CA, Soares EG, et al. Leukotrienes play a role in the control of parasite burden in murine strongyloidiasis. J Immunol. 2005;175: 3892–3899. doi:10.4049/jimmunol.175.6.3892

58. Jiménez M, Cervantes-García D, Córdova-Dávalos LE, Pérez-Rodríguez MJ, Gonzalez-Espinosa C, Salinas E. Responses of Mast Cells to Pathogens: Beneficial and Detrimental Roles. Front Immunol. 2021;12: 685865. doi:10.3389/fimmu.2021.685865

59. Giera M, Kaisar MMM, Derks RJE, Steenvoorden E, Kruize YCM, Hokke CH, et al. The Schistosoma mansoni lipidome: Leads for immunomodulation. Anal Chim Acta. 2018;1037: 107–118. doi:10.1016/j.aca.2017.11.058

60. Thompson-Souza GA, Gropillo I, Neves JS. Cysteinyl Leukotrienes in Eosinophil Biology: Functional Roles and Therapeutic Perspectives in Eosinophilic Disorders. Front Med (Lausanne). 2017;4: 106. doi:10.3389/fmed.2017.00106

61. Bandeira-Melo C, Weller PF. Eosinophils and cysteinyl leukotrienes. Prostaglandins Leukot Essent Fatty Acids. 2003;69: 135–143. doi:10.1016/s0952-3278(03)00074-7

62. Mesquita-Santos FP, Bakker-Abreu I, Luna-Gomes T, Bozza PT, Diaz BL, Bandeira-Melo C. Co-operative signalling through DP(1) and DP(2) prostanoid receptors is required to enhance leukotriene C(4) synthesis induced by prostaglandin D(2) in eosinophils. Br J Pharmacol. 2011;162: 1674–1685. doi:10.1111/j.1476-5381.2010.01086.x

63. Mesquita-Santos FP, Vieira-de-Abreu A, Calheiros AS, Figueiredo IH, Castro-Faria-Neto HC, Weller PF, et al. Cutting edge: prostaglandin D2 enhances leukotriene C4 synthesis by eosinophils during allergic inflammation: synergistic in vivo role of endogenous eotaxin. J Immunol. 2006;176: 1326–1330. doi:10.4049/jimmunol.176.3.1326

64. Amorim NRT, Souza-Almeida G, Luna-Gomes T, Bozza PT, Canetti C, Diaz BL, et al. Leptin Elicits In Vivo Eosinophil Migration and Activation: Key Role of Mast Cell-Derived PGD2. Front Endocrinol (Lausanne). 2020;11: 572113. doi:10.3389/fendo.2020.572113

65. Parente R, Giudice V, Cardamone C, Serio B, Selleri C, Triggiani M. Secretory and Membrane-Associated Biomarkers of Mast Cell Activation and Proliferation. Int J Mol Sci. 2023;24: 7071. doi:10.3390/ijms24087071

66. Bandeira-Melo C, Woods LJ, Phoofolo M, Weller PF. Intracrine cysteinyl leukotriene receptor-mediated signaling of eosinophil vesicular transport-mediated interleukin-4 secretion. J Exp Med. 2002;196: 841–850. doi:10.1084/jem.20020516

67. Neves JS, Radke AL, Weller PF. Cysteinyl leukotrienes acting via granule membrane-expressed receptors elicit secretion from within cell-free human eosinophil granules. J Allergy Clin Immunol. 2010;125: 477–482. doi:10.1016/j.jaci.2009.11.029

68. Zuo S, Wang B, Liu J, Kong D, Cui H, Jia Y, et al. ER-anchored CRTH2 antagonizes collagen biosynthesis and organ fibrosis via binding LARP6. EMBO J. 2021;40: e107403. doi:10.15252/embj.2020107403

69. Bandeira-Melo C, Weller PF, Bozza PT. EicosaCell - an immunofluorescent-based assay to localize newly synthesized eicosanoid lipid mediators at intracellular sites. Methods Mol Biol. 2011;689: 163–181. doi:10.1007/978-1-60761-950-5_10

